# Effect of synchronization of firings of different motor unit types on the force variability in a model of the rat medial gastrocnemius muscle

**DOI:** 10.1101/2020.08.24.264721

**Authors:** Rositsa Raikova, Vessela Krasteva, Piotr Krutki, Hanna Drzymała-Celichowska, Katarzyna Kryściak, Jan Celichowski

## Abstract

Oscillations of muscle force, observed as physiological tremors, rely upon the synchronized firings of active motor units (MUs). This study aimed to investigate the effects of synchronizing the firings of three types of MUs on force development using a mathematical model of the rat medial gastrocnemius muscle. The model was designed based on the actual proportion and physiological properties of MUs and motoneurons innervating the muscle. The isometric muscle and MU forces were simulated by a model predicting non-synchronized firing of a pool of 57 MUs (including eight slow, 23 fast resistant to fatigue, and 26 fast fatigable) to ascertain a maximum excitatory signal when all MUs were recruited into the contraction. The mean firing frequency of each MU depended upon the twitch contraction time, whereas the recruitment order was determined according to increasing forces (the size principle). The synchronization of firings of individual MUs was simulated using four different modes and inducing the synchronization of firings within three time windows (± 2, ± 4, and ± 6 ms) for four different combinations of MUs. The synchronization was estimated using two parameters, the correlation coefficient and the cross-interval synchronization index. The four scenarios of synchronization increased the values of the root-mean-square, range, and maximum force in correlation with the increase of the time window. Greater synchronization index values resulted in higher root-mean-square, range, and maximum of force outcomes for all MU types as well as for the whole muscle output; however, the mean spectral frequency of the forces decreased, whereas the mean force remained nearly unchanged. The range of variability and the root-mean-square of forces were higher for fast MUs than for slow MUs; meanwhile, the relative values of these parameters were highest for slow MUs, indicating their important contribution to muscle tremor, especially during weak contractions.

**Author summary:** The synchronization of firings of motor units (MUs), the smallest functional elements of skeletal muscle increases fluctuations in muscle force, known as physiological tremor, which can disturb high-precision movements. In this study, we adopted a recently proposed muscle model consisting of MUs of three different types (fast fatigable, fast resistant to fatigue, and slow) to study four different scenarios of MU synchronization during a steady level of excitatory input to motoneurons. The discharge patterns were synchronized between pairs of MUs by shifting in time individual pulses, which occurred within a short time interval, and a degree of synchronization was then estimated. The increased synchronization index resulted in increased force variability for all MU types as well as for the whole muscle output; however, the mean force levels remained nearly unchanged, whereas the frequencies of the force oscillations were decreased. The absolute range of force variability was higher for fast than for slow MUs, indicating their dominant influence on muscle tremor at strong contractions, but the highest relative increase in force variability was observed for synchronized slow MUs, indicating their significant contribution to tremor during weak contractions, in which only slow MUs are active.

## Introduction

Most studies of motor unit (MU) firings have revealed the existence of a certain level of synchronization between the firings of motoneurons innervating the same muscle [1-4]. Two concepts for long- and short-term synchronization can be found in the literature. Long-term synchronization with greater latencies beyond ± 20 ms was reported by Datta and Stephens, De Luca et al., Kirkwood et al., Schmied et al., and Semmler et al. [1, 4–7]. The possible mechanism of this kind of synchronization could be explained as interactions occurring between the stretch reflex loop and the recurrent inhibition. Long-term synchronization has been reported to be relatively rare relative to short-term synchronization [4], which was reported to be a peak in the cross-interval histogram centered about a zero-time delay (0.5 ± 2.9 ms). Short-term synchronization is attributed to last-order projections that provide common, nearly simultaneous, excitatory synaptic input across motoneurons [3, 8, 9], generating a narrow peak around the origin of the cross-correlogram of MU discharges [1, 8, 10, 11]. Therefore, the narrow synchronous peak principally reflects shared, monosynaptic projections to motor neurons from corticomotoneuronal cells via the lateral corticospinal tract [12].

In humans, the MU synchronization was shown to be stronger during voluntary muscle activation than during reflex activation [13]. At the same time, synchronization tends to be higher in more distally located muscles, while the greatest synchrony has been most often found in the intrinsic muscles of the foot rather than in the hand muscles [3, 14]. However, the level of synchronization between MUs could be influenced by numerous factors, such as the examined task, the muscles involved in the task, and the type of habitual physical activity performed by the individual [6-7, 15-18]. For example, the level of synchronization was reduced between MUs in the hand muscles of individuals who required greater independent control of the fingers. This included musicians [17] and the dominant hands of control subjects [7]. On the other hand, MU synchronization was found to be greater in the hand muscles of individuals who consistently performed strength training [17, 19] or during tasks that demanded attention [20]. The enhancement of MU synchronization was observed after daily exercise involving brief periods of maximal muscular contraction [19] and contributed to training-induced increments in muscle strength [21]. Better synchronization has also been noted in fatigued muscles [22]. Reports regarding the relationship between physiological tremor and synchronization are inconsistent: most of them have linked tremor with an increased level of synchronization [22-25], while others have suggested no significant associations between the tremor amplitude and the level of MU synchronization exist [17].

It has been assumed that muscle can produce smooth contractions due to asynchronous discharges of motor neurons [23]. Yao et al. [21] revealed that MU synchronization increased the variability in the simulated force but not the average force. Synchronization was also shown to increase the estimated twitch force of the MUs [26].

In the majority of skeletal muscles, three types of MUs have been distinguished and their contractile properties, including the force–frequency of stimulation relationship [27] and sensibility to changes in stimulation pattern [28, 29], were found to vary considerably. In several studies, the effects of the synchronization of MU firings were modelled [21, 30, 31]; however, these models did not analyze the specific effects attributable to different types of MUs. In our previous paper [32], a model of the rat medial gastrocnemius muscle consisting of 30 MUs [10 MUs each of the fast fatigable (FF), fast resistant to fatigue (FR), and slow (S) types] was proposed and the effects of synchronous and asynchronous stimulation of MUs were investigated. It was concluded that the activation of MUs at variable interpulse intervals, delivered to each MU asynchronously, resulted in smaller force oscillations. However, the study did not assess the effects of synchronization between pairs of individual MUs nor the effects of the synchronization of three types of MUs.

A recent model of the rat medial gastrocnemius muscle [33] provided methodology by which to identify the role of each of three MU types (FF, FR, and S) in the production of muscle force. In the present study, the same model was adopted as a tool for simulation of four modes and three time levels of synchronization. The aim of this research was to reveal the important effects of synchronization on the force variability and the force mean spectral frequency and to compare these effects between all types of MUs and the whole muscle. The implication of the results for explanation of tremor at various levels of the muscle force was discussed.

## Materials and methods

### Muscle model

This study applied a model of the rat muscle gastrocnemius based on excitability and firing frequencies of motoneurons, contractile properties, and the number and proportion of MUs in the muscle [33]. Briefly, the model consists of 57 MUs, including eight S, 23 FR, and 26 FF MUs, respectively. As input data, this set of MUs, recorded in physiological experiments, was selected and their twitches were precise modeled by a six-parameter analytical function [34]. The muscle force was calculated as the sum of forces of all active MUs and the process of force regulation was set according to the common-drive hypothesis [35]. The muscle unfused tetanus was calculated following the application of a train of irregular stimuli and was simulated using an analytical approach described in previous research [33, 36]. Meanwhile, the scheme of MU firing was adopted from Fuglevand et al. [30].

In the present study, the excitation signal is simulated (Fig. 1A) as consisting of two smooth logarithmic parts existing during the increasing and decreasing parts of the muscle force (each lasting 1000 ms) and a straight line present during the steady state of the muscle (lasting 2000 ms). The shape of the signal waveform was designed to better approximate more realistically a course of excitation input to motoneurons, avoiding sudden changes occurring in any trapezoidal signal used previously. This study considered only one excitation level, corresponding to 100% of the activation signal, ensuring that all MUs were activated during the steady state of the muscle to enable a thorough analysis of their synchronization. The program for simulation of the force MUs and the muscle force accepted the same MU firing frequencies as previously described (Table 1 in Raikova et al. [33]). The first MU firings at equal interpulse intervals (IPIs) were calculated and, during a second step, a random shifting of IPIs (within intervals of 0, ± 1, and ± 2 ms) was applied, thus simulating a train of firings at irregular IPIs. Finally, the model generated the output forces for different MUs (S, FR, and FF) and the whole muscle, as illustrated in Fig. 1B (sampling frequency *fs* = 1 kHz). This was further denoted as the basic (non-synchronized; *NS*) model, to which no attempts of manual changes of MU firing for synchronization were applied. The force signals were analyzed during the steady-state periods (2000–4000 ms). Their power frequency spectra were calculated by using fast Fourier transform (FFT) over *nf* = 2048 points, thus achieving a spectral resolution Δ*f* = *fs*/*nf* = 0.49 Hz (Figs. 1C–1F). The zero-frequency component defined by the large mean force offset was rejected as soon as it had no relevancy to the frequency components related to the variability of the simulated force, which was under the scope of this study.

**Table 1.** Two indices of synchronization (*corMU* and *CISI*), measured for the *NS* model and four methods of synchronization (Methods 1–4) within three time windows (± 2, ± 4, and ± 6 ms). All values are reported as the mean value ± standard deviation for different physiological types of MUs (S, FR, and FF) and for all MUs within the muscle.

|  |  | **corMU [%]** |  |  |  | **CISI [%]** |  |  |  |
| --- | --- | --- | --- | --- | --- | --- | --- | --- | --- |
|  |  | **S MUs** | **FR MUs** | **FF MUs** | **all MUs** | **S MUs** | **FR MUs** | **FF MUs** | **all MUs** |
| **NS** |  | **4.5±2.0** | **7.4±2.9** | **5.7±2.8** | **6.1±2.8** | **5.8±0.5** | **6.2±0.4** | **6.2±0.3** | **6.2±0.4** |
| **Method 1** | **±2ms** | 10.4±7.8 | 11.5±8.6 | 8.2±7.6 | 7.3±5.8 | 6.4±0.8 | 7.8±0.6 | 7.3±0.7 | 7.4±0.8 |
|  | **±4ms** | 21.9±15.2 | 21.3±16.5 | 13.4±14.7 | 10.2±11.4 | 8.1±0.9 | 11.7±0.9 | 9.6±1.6 | 10.3±1.8 |
|  | **±6ms** | 37.2±19.5 | 38.4±21.1 | 22.3±22.4 | 15.0±18.4 | 8.8±1.0 | 18.7±1.8 | 13.9±3.4 | 15.1±4.3 |
| **Method 2** | **±2ms** | 10.4±9.7 | 11.4±10.6 | 8.4±9.0 | 7.6±6.6 | 6.7±0.5 | 8.0±1.0 | 7.6±0.8 | 7.6±0.9 |
|  | **±4ms** | 20.6±15.7 | 22.5±19.0 | 13.9±15.9 | 10.5±12.5 | 7.6±0.5 | 12.3±1.8 | 10.0±1.8 | 10.6±2.3 |
|  | **±6ms** | 38.1±19.7 | 39.7±21.5 | 20.2±21.9 | 15.2±18.4 | 10.3±0.8 | 19.4±3.1 | 13.3±2.9 | 15.4±4.5 |
| **Method 3** | **±2ms** | 9.5±7.7 | 10.0±7.9 | 7.5±7.5 | 6.9±5.4 | 6.4±0.6 | 7.2±0.4 | 6.9±0.7 | 6.9±0.6 |
|  | **±4ms** | 15.7±14.8 | 13.5±16.2 | 9.9±13.9 | 7.9±9.9 | 7.2±0.8 | 8.4±0.5 | 7.9±1.2 | 8.0±1.0 |
|  | **±6ms** | 23.8±23.5 | 16.8±24.3 | 12.2±20.0 | 9.1±14.5 | 8.2±1.0 | 9.7±0.7 | 9.0±1.4 | 9.2±1.2 |
| **Method 4** | **±2ms** | 11.5±6.9 | 19.5±7.6 | 16.6±9.3 | 10.1±8.2 | 6.3±0.7 | 10.8±1.6 | 10.9±1.9 | 10.2±2.3 |
|  | **±4ms** | 24.9±10.6 | 46.6±9.6 | 39.3±13.5 | 19.1±19.5 | 7.8±1.1 | 21.4±2.1 | 21.0±2.4 | 19.3±5.2 |
|  | **±6ms** | 42.6±10.9 | 74.6±6.9 | 62.8±13.1 | 28.3±31.2 | 9.0±1.4 | 32.1±1.5 | 31.6±1.4 | 28.6±8.1 |

**Figure 1.**
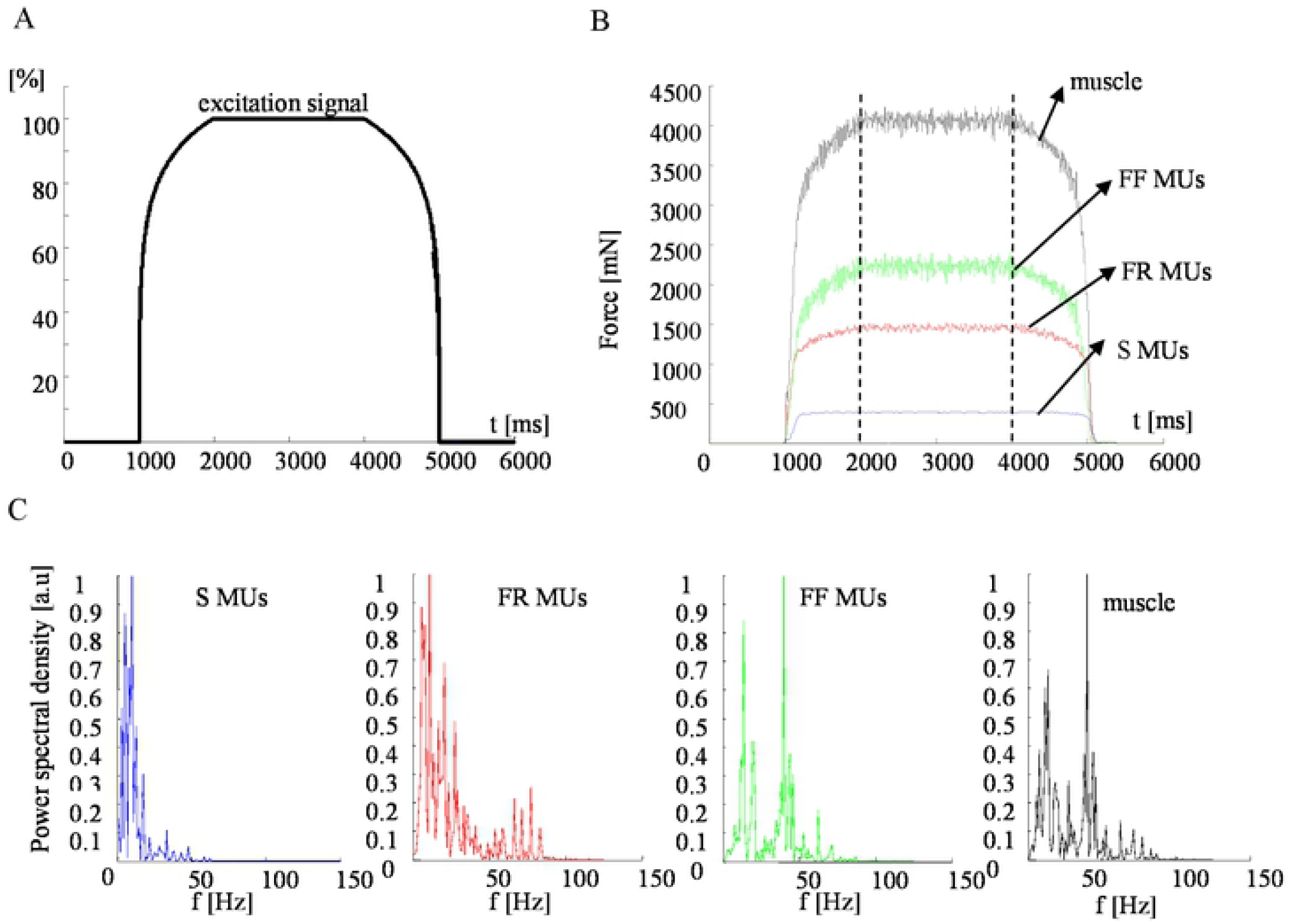
Parameters of the basic model, calculated using a 100% excitation signal. A. The law for the excitation. B. The calculated forces of populations of different MU types (S, FR, and FF) and the muscle. C. Normalized power spectral density of the force during a time period of 2000 to 4000 ms, presented separately for individual MUs (S, FR, and FF) and the whole muscle.

### Simulation of MU synchronization

The *NS* firings of all 57 MUs during the muscle steady state are shown in Fig. 2. These patterns were further modified to simulate different types and levels of synchronization. The synchronization was applied to a specific pair of MUs (named MU1 and MU2) so that the impulses of MU2, which fall within a predefined time window, Δt, around the impulses of MU1, were changed to coincide with those of MU1. Three time windows with Δt = ± 2, ± 4 or ± 6 ms were used to simulate three levels of MU synchronization, mimicking weak, modest, and strong synchronization, respectively. The synchronization scheme is illustrated in Fig. 3, showing that the larger the time window was, the greater number of more MU pulses were shifted to and synchronized with the reference MU.

**Figure 2.**
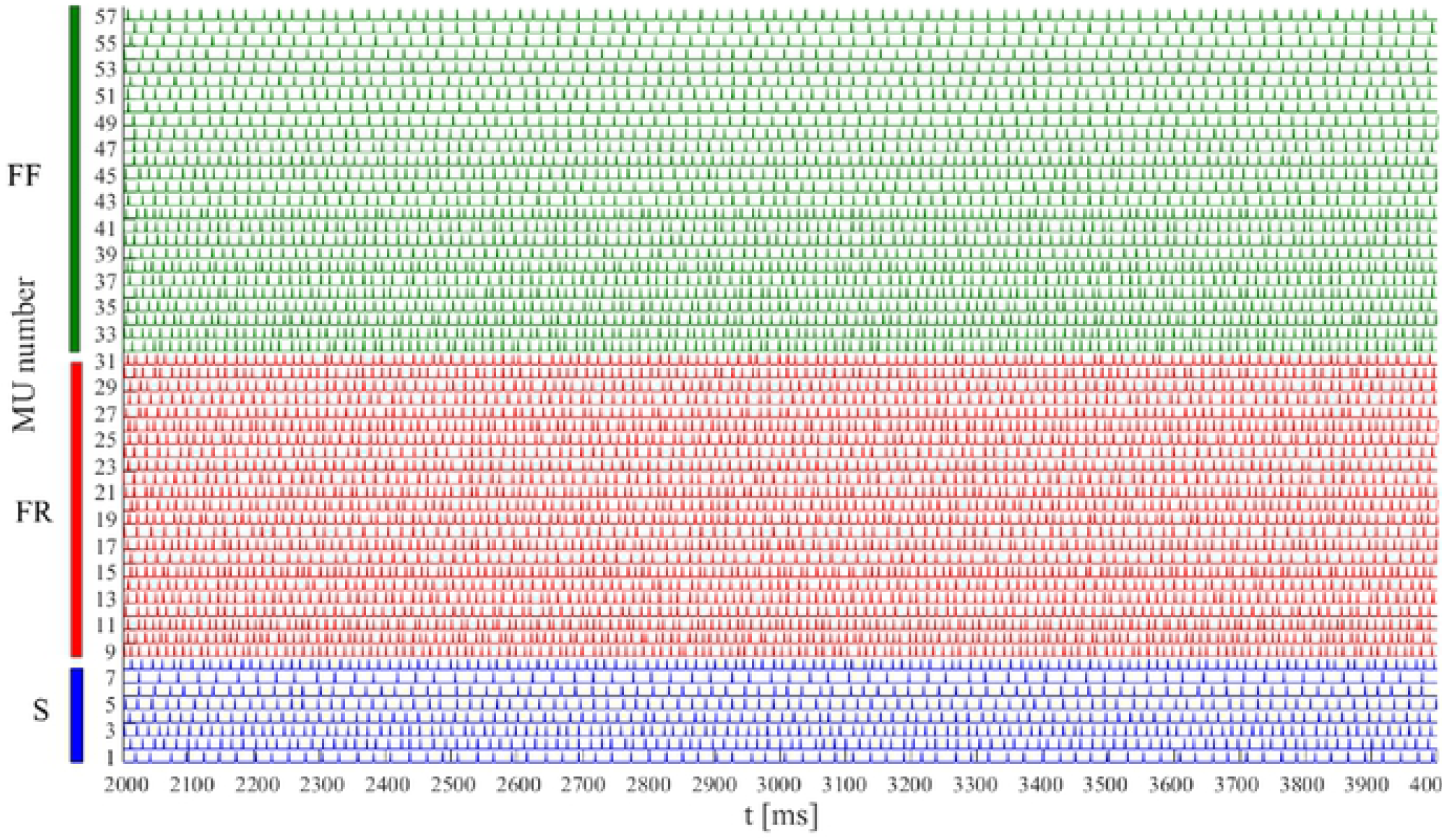
Firing patterns of 57 MU of the basic *NS* model during the time period of 2000 to 4000 ms. MUs are numbered in an ascending order based on their maximum twitch forces within each type S (1–8), FR (9–31), and FF (32–57).

**Figure 3.**
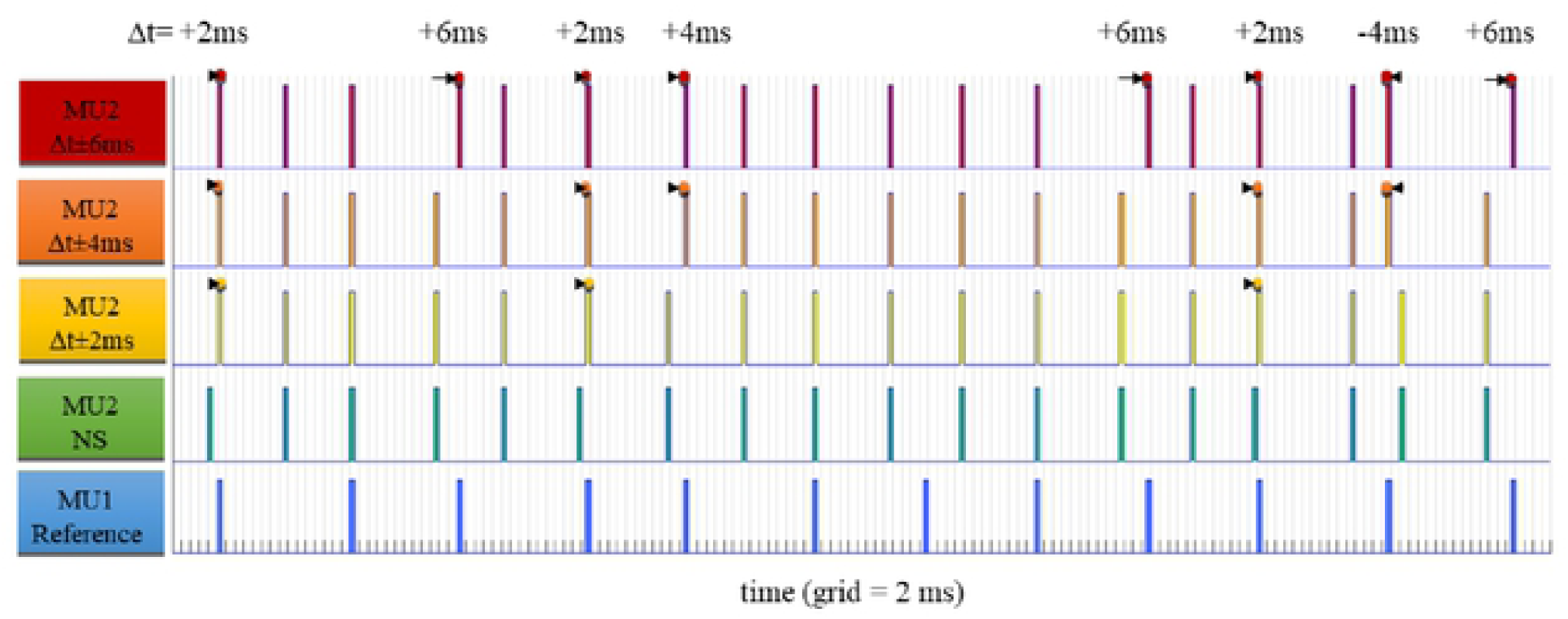
Illustration of the synchronization principle of basic MU firing patterns of two MUs, considering MU1 as a reference one (blue, bottom) and applying the synchronization of pulses to MU2 (green, *NS*). Three time windows were used: Δt = ± 2 ms (yellow), ± 4 ms (orange), and ± 6 ms (red). The dots highlight individual pulses ofMU2, which were shifted in time (left or right, as indicated by arrows) to coincide with reference impulses of MU1 when the time interval between the pair of impulses of MU2 and MU1 was less or equal than |Δt|. The level of synchronization was proportional to Δt, illustrated in this example by increasing numbers *n* of shifted impulses-namely, *n* = 3 for Δt = ± 2 ms, *n* = *5* for Δt = ± 4 ms, and *n* = 8 for Δt = ± Δ ms.

Four methods of synchronization (Methods 1–4) were applied. In all methods, the synchronized MU pairs were chosen only encompassing the same physiological type (S and S, FR and FR, FF and FF), i.e., synchronization was not induced between MUs of different types.

*Method 1*: Two neighboring MUs within the same physiological type according to the recruitment order based on their increasing force of the twitch (see Table 1 in Raikova et al. [33]) were synchronized, i.e., for S MUs, 1–2, 2′–3, …, 7′–8; for FR MUs, 9–10, 10′–11, …, 30′–31; and, for FF MUs, 32–33, 33′–34, …, 56′–57. Note that, for each next synchronization, the already synchronized pattern of the previous MU is used and marked by “′”.

*Method 2*: Two neighboring MUs within the same physiological type but when ordered according to their increasing mean firing rate (see Table 1 in Raikova et al. [33]), were synchronized i.e., for S MUs, 7-1, 1′–6, 6′-5, 5′–4, 4′–2, 2′–3, and 3′–8; for FR MUs, 18–16, 16′–24, 24′–22, 22′–28, 28′–14, 14′–12, 12′–23, 23′–13, 13′–20, 20′–31, 31′–27, 27′–29, 29′– 25, 25′–9, 9′–21, 21′–30, 30′–26, 26′–19, 19′–17, 17′–11, 11′–10, and 10′–15; and, for FF MUs, 50–44, 44′–43, 43′–49, 49′–39, 39′–54, 54′–52, 52′–55, 55′–56, 56′–48, 48′–47, 47′– 53, 53′–51, 51′–37, 37′–40, 40′–41, 41′–57, 57′–45, 45′–35, 35′–33, 33′–46, 46′–32, 32′–34, 34′–38, 38′–36, and 36′–42.

*Method 3*: The MUs within the same physiological type but in unique groups of four MUs were synchronized to the first recruited MU and ordered according to their increasing force of the twitch (see Table 1 in Raikova et al. [33]), i.e., for S MUs, 1–2, 1–3, 1–4, 5–6, 5–7, and 5–8; for FR MUs: 9–10, 9–11, 9–12, 13–14, 13–15, 13–16, 17–18, 17–19, 17–20, 21–22, 21– 23, 21–24, 25–26, 25–27, 25–28, 29–30*, and 29–31*; and, for FF MUs, 32–33, 32–34, 32– 35, 36–37, 36–38, 36–39, 40–41, 40–42, 40–43, 44–45, 44–46, 44–47, 48–49, 48–50, 48–51, 52–53, 52–54, 52–55, and 56–57*. The symbol (*) denotes the groups, where the number of synchronized MUs was less than four due to the fact that the number of MUs in the respective physiological type was not a multiple of four.

*Method 4*: The MUs within the same physiological type were synchronized, taking as a reference the first recruited MU of the specific type (see Table 1 in Raikova et al., [33]), i.e., for S MUs, 1–2, 1–3, …, 1–8; for FR MUs, 9–10, 9–11, …, 9–31; and, for FF MUs, 32–33, 32–34, …, 32–57.

### Estimation of MU synchronization

#### Temporal correlation of MU impulses

The MU pulses were represented as an MU binary (MUB) sample series with a constant sampling period of 1 ms and binary amplitude of 0 or 1, where “0” indicated a non-active state and “1” indicated the presence of a pulse-active state. The duration of the pulse-active state was set to 1 ms, overlaying one sampling period. MUB series were represented with a total of 2000 samples during the steady state of the muscle from 2000 ms to 4000 ms, as depicted in Figs. 1 and 2 for the MUs in the basic, *NS* model.

The temporal correlation between the binary sample series of two MUs (MUB1 and MUB2) was computed with the normalized Pearson’s correlation coefficient ranged in the interval 0% to 100%, according to the following formula:

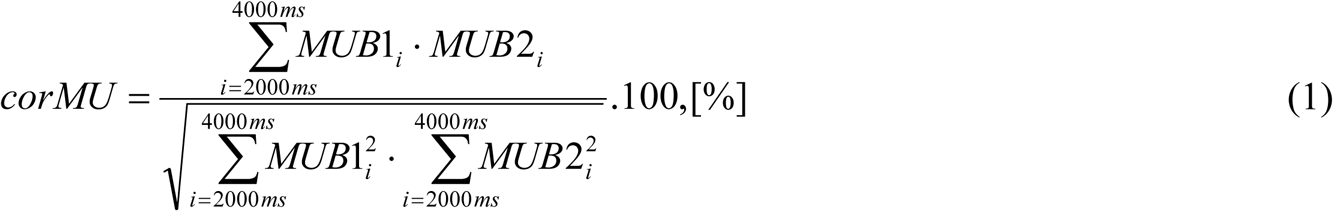

where *i* denotes the sample index of the MUB series, considering a sampling period of 1 ms.

The correlation coefficient (*corMU*) is a standard measure of similarity between sample series data in the time domain. Transferring this knowledge to the MUB data, *corMU* is representative of the temporal synchronization of two MU firings such that 100% corresponds to a complete coincidence between all firing pulses in MU1 and MU2 and 0% corresponds to no coincidence between any firing pulse in MU1 and MU2. The normalized value of *corMU* does not depend upon the length of the estimated MUB time series, the number of firing pulses, or the mean firing rate,. This is an important benefit of the normalization, which would prevent bias in *corMU* estimation, considering that MUs in different physiological types have different mean firing rates.

#### Cross-interval synchronization index

The synchronization between the firing patterns of two MUs (MU1 and MU2) was estimated by an analysis of their cross-intervals using *CI*_*x*_ (*MU*1, *MU* 2) = {*t*1_*x*_ − *t*2_*xy*_} computed as a pair-wise difference between the times of occurrence of all reference MU1 pulses *t*1_*x*_ = {*t*1_1_, *t*1_2_, ….*t*1_*nMU* 1_} and their corresponding closest neighbors among MU2 pulses *t*2_*xy*_ ∈ *t*2 _*y*_ = {*t*2_1_, *t*2_2_, *t*2_*nMU* 2_}. The latter were found by the minimization criterion 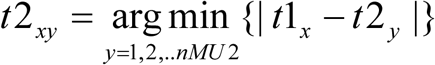 and respected the condition to overlay only firings during the steady state of the muscle, i.e., *t*1_*x*_, *t*2_*y*_ ∈ [ 2000*ms;* 4000*ms]*. By definition, the *CI* _*x*_ (*MU* 1, *MU* 2) vector length was equal to the number of pulses in the reference MU (*nMU1*). *CI* values could be negative, zero, or positive when an MU1 pulse was respectively preceding, coinciding with or following its neighbor MU2 pulse, as illustrated in Fig. 4.

**Figure 4.**
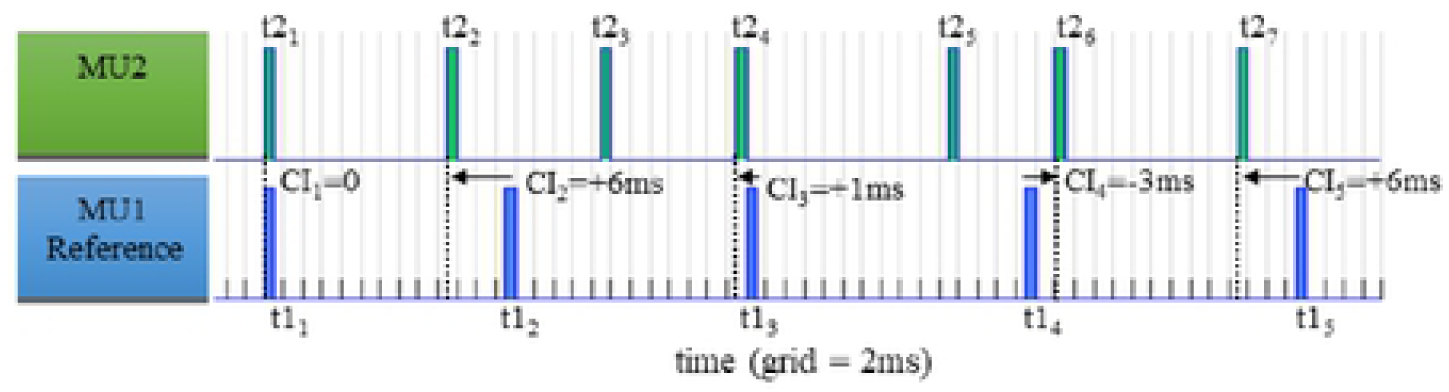
Illustration of the cross-interval measurements of pulses of real MU firing patterns {CI_1_, CI_2_, CI_3_, CI_4_, CI_5_}, considering MU1 as a reference (blue, bottom) and applying pairwise differences between the times of occurrences of all MU1 pulses {t1_1_, t1_2,_ t1_3_, t1_4_, t1_5_} and their corresponding closest neighboring pulse times of MU2 {t2_1_, t2_2_, t2_4_, t2_6_, t2_7_}.

The distribution of cross-interval values of two MUs, *CI(MU1,MU2)*, was estimated by means of a cross-interval histogram with a bin-width resolution of 1 ms and bin centers in the range of ± 15 ms. The bin values represented the relative probability (*p*_*bi*_) of having a *CI* observation within a specific bin interval (*bi):*

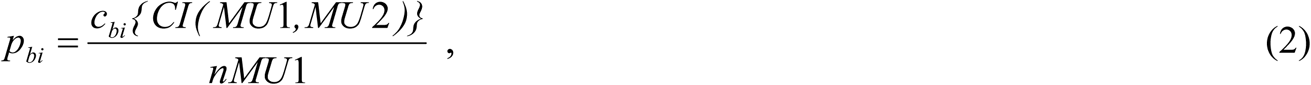

accepting the sum of all bin values equal to 1:

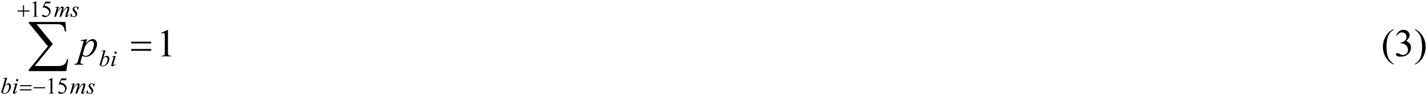

where *c*_*bi*_ is the count of *CI (MU1,MU2)* values in bin *bi* and the denominator is the number of elements in the input data, equal to the number of reference MU pulses (*nMU1*).

Derived from the cross-interval histogram, the synchronization between the firing patterns of MU1 and MU2 was estimated by the relative probability *p*_*b0*_ in the central bin (*b0* = ± 0.5 ms), equivalent to the relative frequency of coincidence between MU1 and MU2 pulses related to the reference number of pulses:

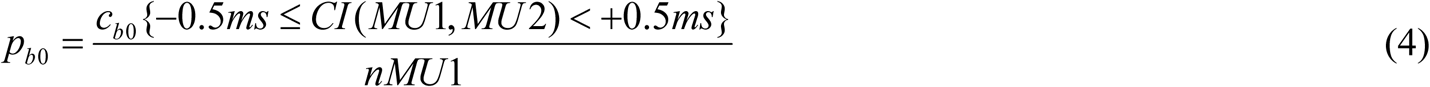

Given a total number of N = 57 MUs, there could be derived a total of N-1 cross-interval vectors *CI(MUi, MUj)* for any given pair of MUs, where *i, j* = 1, 2,.., N, and i≠j. Further, a cross-interval synchronization index (*CISI*) was defined for each reference MUi pattern to accumulate the relative probability of pulse coincidences in all MUi pairs (*N* − 1) observed in the central bin:

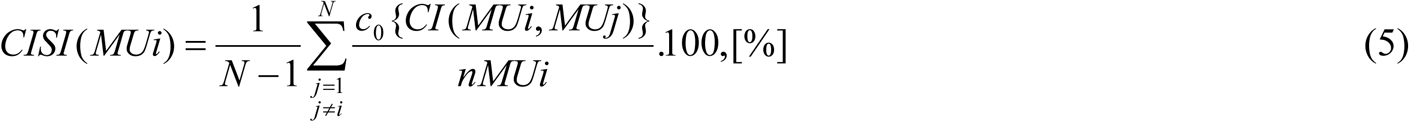

*CISI* has a normalized value from 0 to 100% with 0% corresponding to no coincidence and 100% corresponding to a complete coincidence between the patterns of the reference MUi and all other paired MUs. The adopted *CISI* normalization to both number of MU pairs (*N*) and number of reference firings (*nMUi*) was implemented to reject the influence of the size and type of the studied MU population.

#### Force parameters

Standard force parameters of the different MU groups (S, FR, and FF) and the cumulative force of the whole muscle (Fig. 1B) were calculated as follows:

- Force mean value: 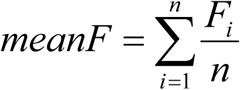
- Force max value: *max F* = *max(F*_*i*_ )
- Force min–max range: *rangeF* = *max(F*_*i*_ ) − *min(F*_*i*_ )
- Force root-mean-square (*RMS*) level: 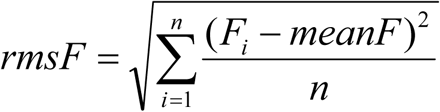

where *F*_*i*_ denoted the force signal samples, taken with a sampling period of 1 ms during the steady state of the muscle from 2000 ms to 4000 ms, including a total number of *n* = 2000 samples.

Additionally, the force power spectral density (*PSD*) of different MU groups (S, FR, and FF) and the cumulative muscle force (Fig. 1C) was used for the calculation of the mean spectral frequency as follows:

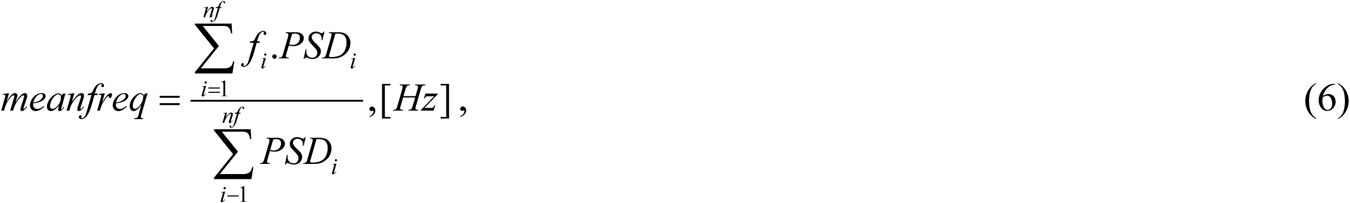

where *nf* is number of frequency bins in the spectrum (*nf* = 2048 as defined earlier), *fi* is the frequency of the spectrum at bin *i* of *nf*, and *PSDi* is the amplitude of the *PSD* at bin *i* of *nf*.

## Results

### Weak synchronization of MU firings in the basic muscle model

The level of synchronization between MU firing patterns of the simulated basic rat muscle gastrocnemius model with 100% excitation containment and 57 MUs, over a two -second time period during the muscle steady state, was estimated by the two different synchronization indices in Table 1 (top row) and discussed as follows.

First, *corMU* = 6.1% ± 2.8% (mean value ± standard deviation) shows a ***weak temporal*** correlation between the firing pulses of all MUs within the muscle, which was found to be lowest for MUs of type S (4.5% ± 2%) and highest for those of type FR (7.4% ± 2.9%). A comprehensive proof for the absence of noteworthy clusters with significant correlation between MUs of a specific type is illustrated in the *corMU* colormap in Fig. 5A. Here, a random distribution of *corMU* values can be noted, overlaying the dark-blue colored area of very low pairwise correlations between 57 × 57 MUs, distributed on the x- and y-axes. The entries in the main diagonal should be ignored because each represents a MU compared with itself (*corMU* = 100%).

**Figure 5.**
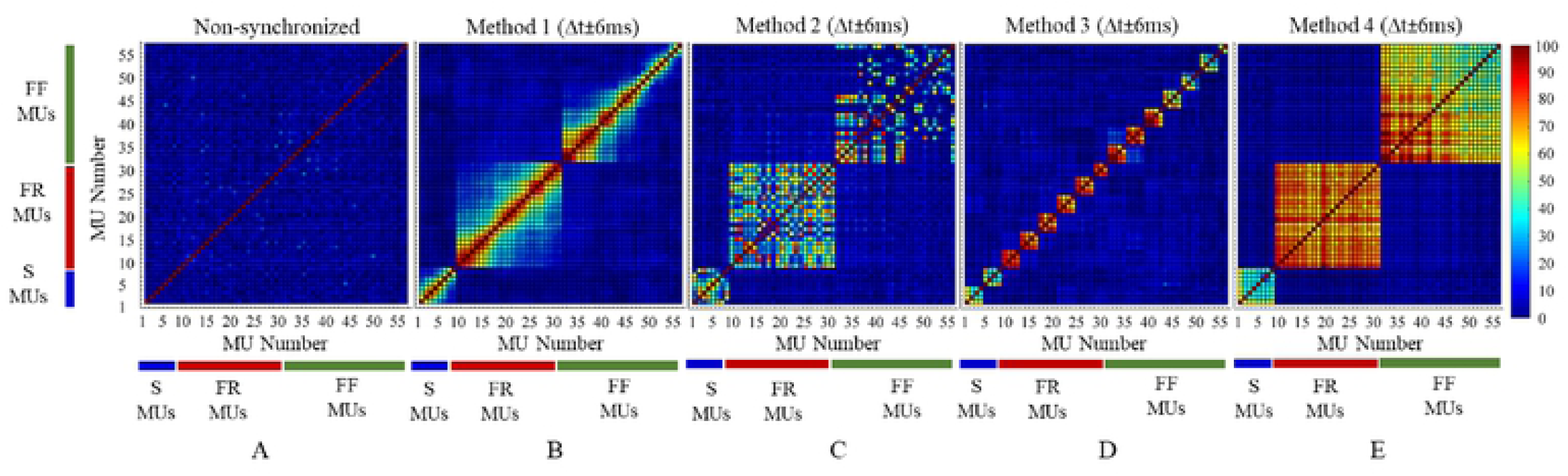
The correlation coefficients (*corMU*) between all pairs of 57 MU firing patterns for the *NS* model and for the four methods of MU synchronization using Δt ± 6 ms. The color map represents *corMU* values in the range of 0% to 100% calculated within the square grid (57 ⨯ 57) of sequential MU numbers from 1 to 57. The diagonal elements of the color map correspond to a 100% correlation between the firing pulses of identical MUs.

Second, *CISI* = 6.2% ± 0.4% (mean value ± standard deviation) suggests ***weak*** cross-interval synchronization between the firing pulses of all MUs within the muscle, without essential differences between MUs of different physiological types [the *CISI* mean value varied from 5.8% (S MUs) to 6.2% (FR and FF MUs)]. Evidence for missing synchronization between MU firings can be observed in the cross-interval histograms in Fig. 6A, having a flat (uniform) distribution in the range of bin-intervals [−6 ms; +6 ms] for all 57 MUs. Therefore, cross-intervals between firing patterns were equally probable within this bin range and no evidence for synchronous peaks could be identified in the case of any MU.

**Figure 6.**
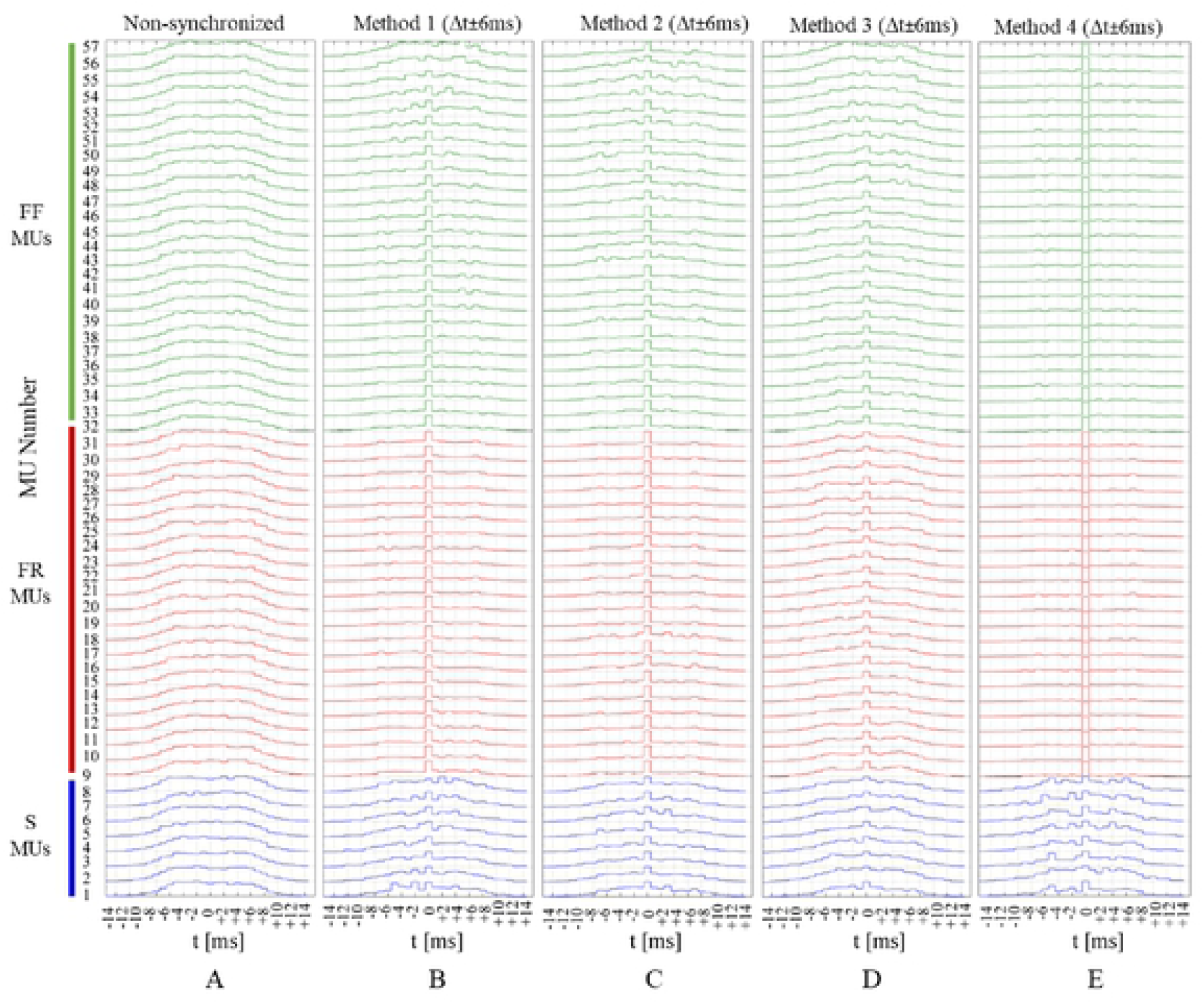
Cross interval histograms of all 57 MU firing pattems for the *NS* basic model and the four methods of MU synchronization using Δt ± 6 ms. The cross-interval histograms are depicted with maximal bin nonnalization, considering a bins width of 1 ms within a bin interval of ± 15 ms. The amplitude of the central bin, presenting a minimal cross-interval difference of ± 0.5 ms, is proportional to the derived index of synchronization (*CISI*).

### Stronger synchronization of MU firings in different synchronization scenarios

The aforementioned 57 MU firing patterns of the basic muscle model were modified according to 12 synchronization scenarios, i.e., four synchronization concepts (Methods 1–4) each applied within three synchronization time intervals (Δt = ± 2, ± 4, and ± 6 ms). The resultant average levels of synchronization between patterns of MUs of the same physiological type and within the whole muscle are estimated in Table 1. In all cases, certain increments of both indices for the level of MU synchronization (*corMU* and *CISI*) were assessed in comparison with their estimation for the basic *NS* model in the first row of Table 1. Therefore, it may be concluded that the simulation design achieved the general goal for inducing stronger synchronization between MU firings. More details on the observed MU synchronizations related to the computation of *corMU* and *CISI* are presented below.

- *corMU*: Different effects of the synchronization induced by Methods 1 tough 4 could be tracked well on the *corMU* color map (Figs. 5B–5E), seen as clusters with strong correlations (*corMU* is from 30% to 100%). These clusters have different two-dimensional space distributions of the entries with maximal correlation, corresponding to the different concepts for synchronization of MU pairs in Methods 1 through 4, as follows:
  - *Method 1*: The synchronization between neighboring MUs is seen in Fig. 5B as maximal correlations around the main diagonal (identical MUs and their closer neighbors) and a trend of gradually decreasing correlations moving away from that diagonal (MU pairs with far neighborhood). Three clusters with *corMU* gradient can be identified in Fig. 5B as a result of synchronization within MUs of the same physiological type (S–S, FR–FR, FF–FF). Within these clusters, the maximal correlation (mean value ± standard deviation) is observed for FR MUs (38.4% ± 21.1%), S MUs (37.2% ± 19.5%), and minimally for FF MUs (22.3% ± 22.4%), considering the setting with a maximal synchronization interval Δt = ± 6 ms (Table 1). This means up to 30% increase in the correlation coefficients within MU groups, as compared with in the basic *NS* model.
  - *Method 2*: The synchronization was applied to not ordered MU pairs within the same physiological type; therefore, the *corMU* color map in Fig. 5C appears with a non ordered colorful distribution with strong correlations between various MU pairs, forming three clusters within MUs from the same physiological type (S–S, FR–FR, FF–FF). Within these clusters, maximal correlation (mean value ± standard deviation) was observed for FR MUs (39.7% ± 21.5%), S MUs (38.1% ± 19.7%), and minimally for FF MUs (20.2% ± 21.9%), considering the setting with a maximal synchronization interval Δt = ± 6 ms (Table 1). We note that the reported average *corMU* values in Method 2 are very similar to those in Method 1. Considering that both methods had a common concept for MU synchronization in pairs, we could deduce that the synchronization concept and not the order of MU recruitment can help in increasing the synchronization index by up to 30%, although the effect on the output force is expected to be different.
  - *Method 3*: The synchronization between unique groups of four neighboring MUs is seen in Fig. 5D as maximal correlations in clusters with (4 × 4) entries, distributed around the main diagonal (including the identical MU pair and its three closest neighbors). There are two exceptions with smaller clusters, including 3 × 3 entries (MU numbers 29, 30, 31) and 2 × 2 entries (MU numbers 56, 57), which exactly correspond to the methodological constraints. Considering all MUs within the same physiological type, the maximal correlation (mean value ± standard deviation) is estimated for S MUs (15.7% ± 14.8%), FR MUs (13.5% ± 16.2%), and minimally for FF MUs (9.9% ± 13.9%) in the setting with a maximal synchronization interval Δt = ± 6 ms (Table 1). This result yields an increment of 6% to 18% of *corMU* after Method 3 synchronization relative to with the basic *NS* model. In general, Method 3 induces a smaller level of synchronization than Methods 1 and 2, which can be deduced from the larger size of the dark blue color area with uncorrelated MU pairs found in Fig. 5D as compared with in Figs. 5B and 5C.
  - *Method 4*: The concept for synchronization of all MUs within the same physiological type to only one reference MU resulted in MU clusters with very high pairwise correlations, enclosing all MUs in the respective physiological type (S–S, FR–FR, FF–FF). Within these clusters, the maximal correlation (mean value ± standard deviation) was observed for FR MUs (74.6% ± 6.9%), FF MUs (62.8% ± 13.1%), and minimally for S MUs (42.6% ± 10.9%) in the setting with a maximal synchronization interval Δt = ± 6 ms (Table 1). This result yields an increment from 37% to 67% of the correlations within MU groups relative to with the basic *NS* model and can be denoted as the maximal synchronization level simulated in this study.
- *CISI*: The effect of synchronization induced by Methods 1 through 4 could be identified in the cross-interval histograms (Figs. 6B–6E) by the prominent peak in the central bin (± 0.5 ms). The larger is amplitude deviation from the uniform distribution in other bins, the higher the probability for synchronization of the respective MU to the firing pulses of other MUs. Different synchronization methods produce different amplitudes in the central bin, estimated by *CISI* in Table 1. Comparing the *CISI* values of all methods estimated with maximal synchronization interval Δt = ± 6 ms, we could deduce the following:
  - The lowest *CISI* mean value was found for S MUs (from 8.2% in Method 3 to 10.3% in Method 2, with the latter being up to 4.5% above the basic *NS* model).
  - The largest *CISI* mean value was found for both FF MUs (from 9% in Method 3 to 31.6% in Method 4) and FR MUs (from 9.7% in Method 3 to 32.1% in Method 4). Thus, the best synchronization of Method 4 achieved up to a 25.9% greater *CISI* value as compared with the basic *NS* model.

Additionally, Fig. 8A was designed to show the effect of widening the time window for synchronization (Δt = ± 2, ± 4, and ± 6 ms) on the relative *CISI* change (ratio of synchronized vs. *NS* value). It shows that, generally, Δt = ± 2 ms leads to weak synchronization and slight increases in *CISI* by about 1.1 times (Methods 1–3) and 1.8 times (Method 4); Δt = ± 4 ms lead to maximal synchronization that is still less than two times (Methods 1–3) but about three times (Method 4); and Δt = ± 6 ms produced the maximal synchronization with notable *CISI* increment increases by up to three times (Methods 1 and 2) and up to five times (Method 4).

**Figure 7.**
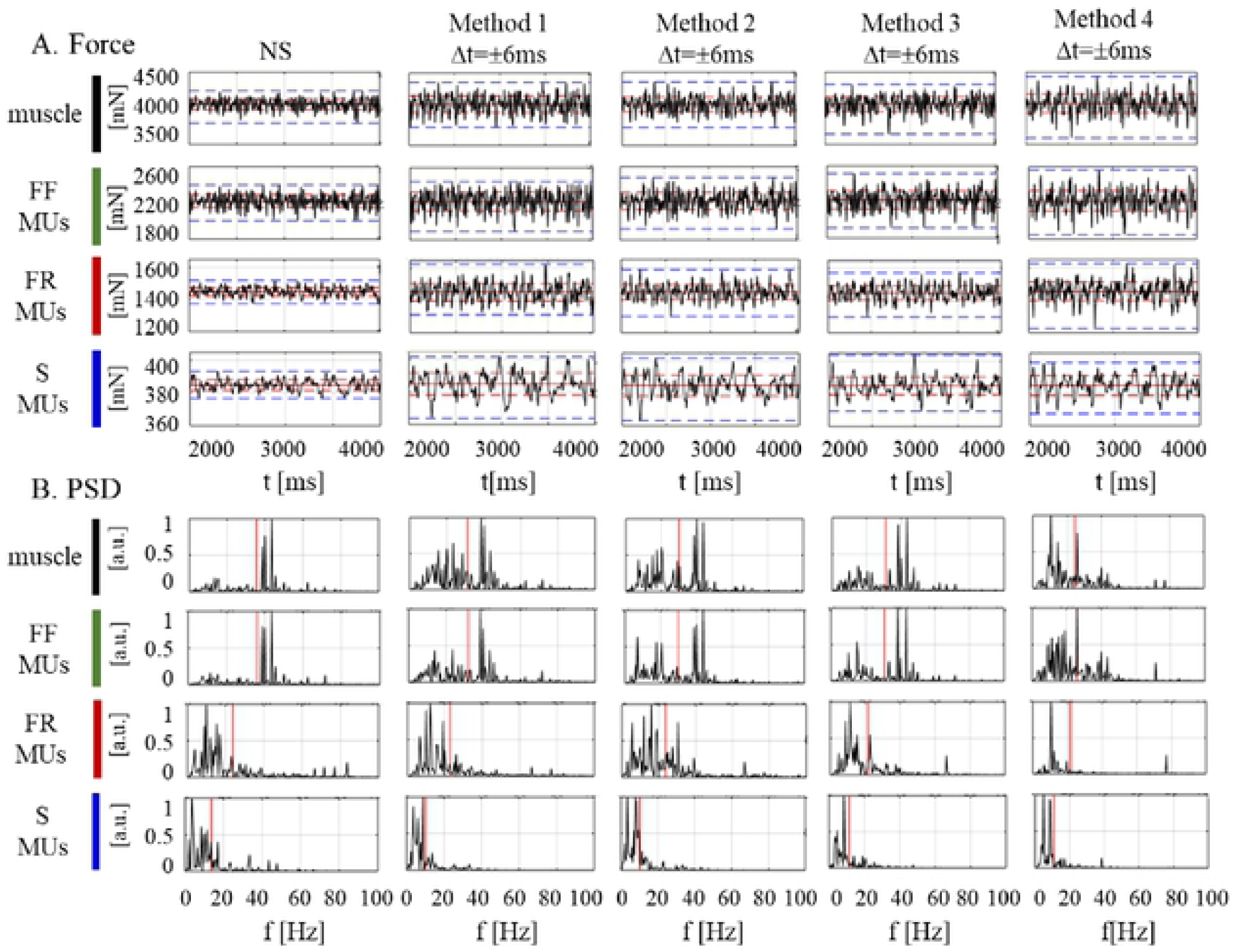
The forces (A) and respective power spectral densities (B) calculated for the muscle and different MU types (S, FR, and FF) during the muscle steady state of the *NS* excitation and by using four methods of MU synchronization (Δt ± 6 ms). Values of different force parameters are indicated in each box of panel A as follows: the mean force by a red horizontal solid line, the force *rms* by a red horizontal dotted line, and the force range by a blue horizontal dotted line; meanwhile, the mean frequency is indicated in each box of panel B by a red ve1tical tick line.

**Figure 8.**
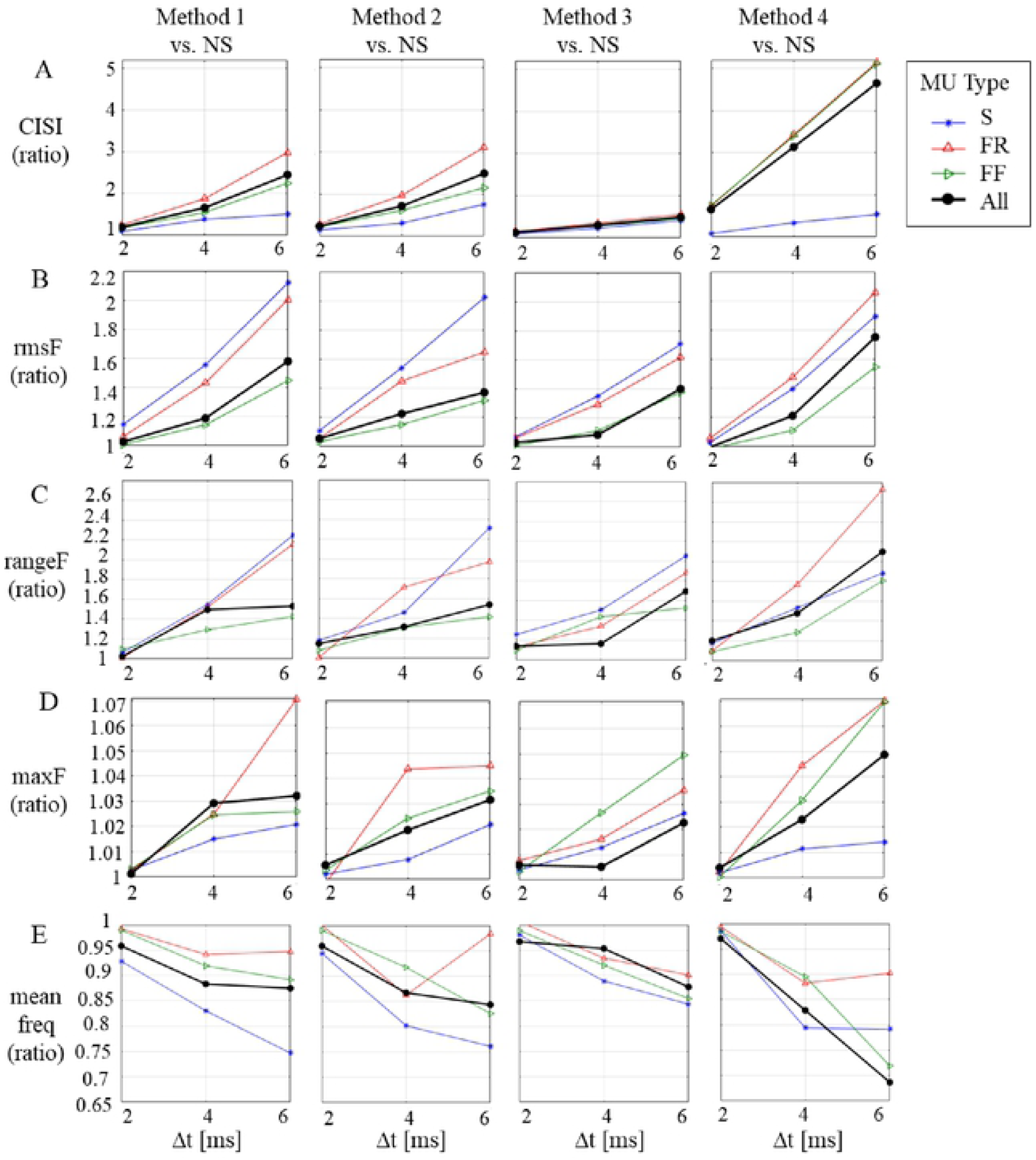
Effects of widening the time window for synchronization (Δt = ± 2, ± 4, and ± 6 ms) on an increment of the cross-interval synchronization index of MU pulses (CISI in panel A), the force *rms* (panel B), the force range (panel C), the maximal force (panel D), and the force mean spectral frequency (panel E), presented as the ratio of values calculated for each method of synchronization (Methods 1–4) vs. the *NS* model.

### Maximal effect of MU synchronization on the force parameters

The forces produced by the muscle and different MU types before and after the application of different synchronization scenarios were estimated for a two -second period during the muscle steady state and the defined five basic force parameters (*meanF, rmsF, rangeF, maxF*, and *meanfreq*) are presented in Table 2. For comprehension purposes, the representation of those parameters on the force signals and their PSD is additionally illustrated in Fig. 7. The comparison of the *NS* excitation to those achieved with different synchronization methods (Methods 1–4) is presented below.

**Table 2.**
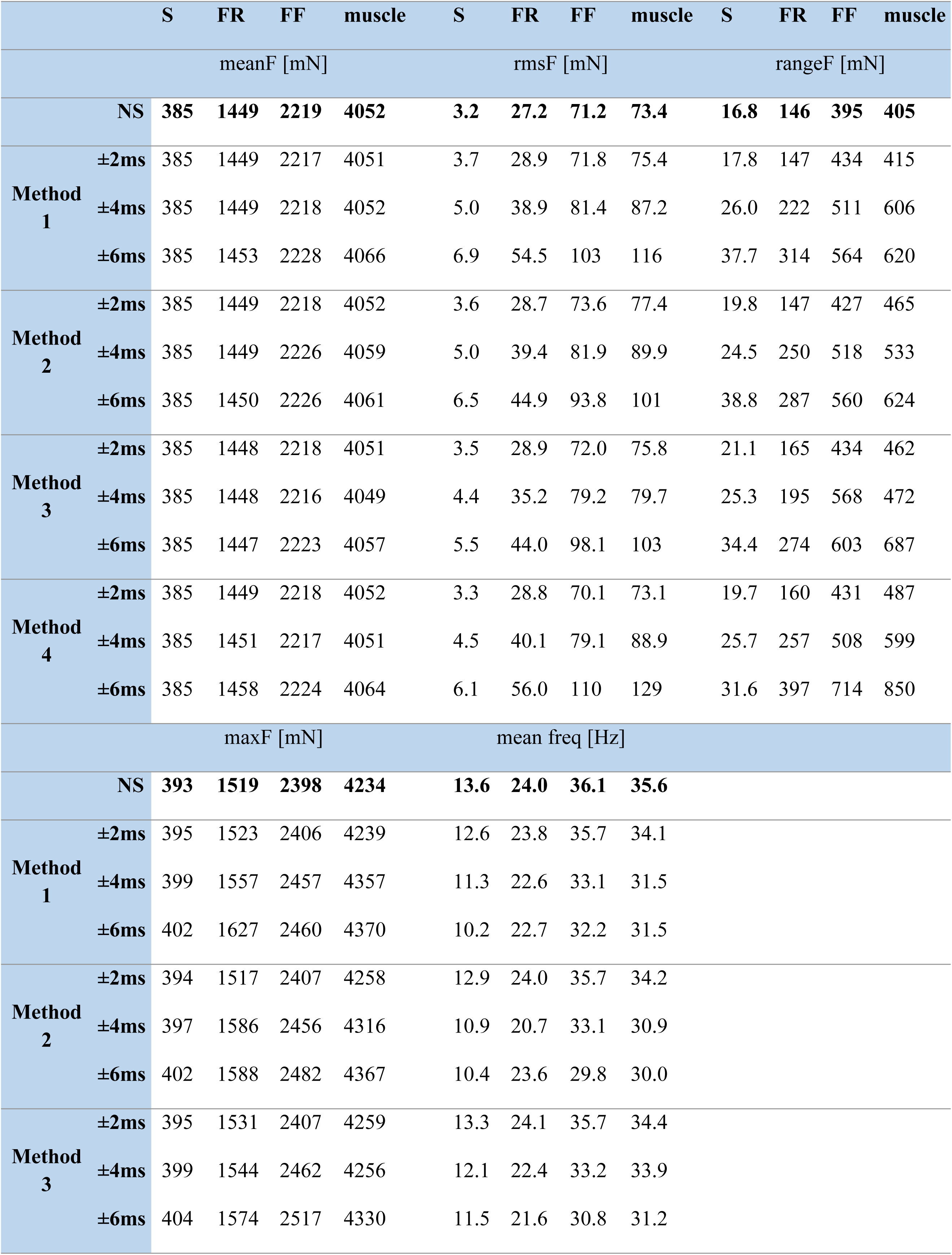
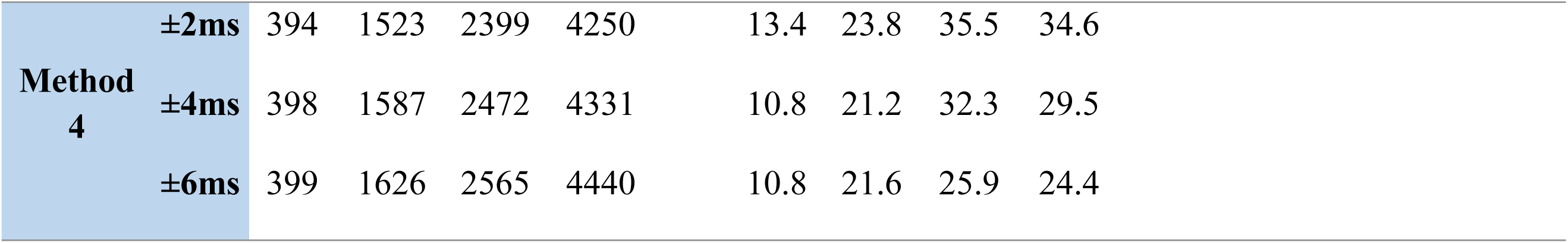
Estimation of the five force parameters *meanF, rmsF, rangeF, maxF*, and *meanfreq* of different physiologicl types of MUs (S, FR, and FF) and the whole muscle, produced by the *NS* model and four methods for synchronization (Methods 1–4), within three time windows (± 2, ± 4, and ± 6 ms).

|  |  | S | FR | FF | muscle | S | FR | FF | muscle | S | FR | FF | muscle |
| --- | --- | --- | --- | --- | --- | --- | --- | --- | --- | --- | --- | --- | --- |
|  |  | meanF [mN] |  |  |  | rmsF [mN] |  |  |  | rangeF [mN] |  |  |  |
|  | NS | 385 | 1449 | 2219 | 4052 | 3.2 | 27.2 | 71.2 | 73.4 | 16.8 | 146 | 395 | 405 |
| Method 1 | ±2ms | 385 | 1449 | 2217 | 4051 | 3.7 | 28.9 | 71.8 | 75.4 | 17.8 | 147 | 434 | 415 |
|  | ±4ms | 385 | 1449 | 2218 | 4052 | 5.0 | 38.9 | 81.4 | 87.2 | 26.0 | 222 | 511 | 606 |
|  | ±6ms | 385 | 1453 | 2228 | 4066 | 6.9 | 54.5 | 103 | 116 | 37.7 | 314 | 564 | 620 |
| Method 2 | ±2ms | 385 | 1449 | 2218 | 4052 | 3.6 | 28.7 | 73.6 | 77.4 | 19.8 | 147 | 427 | 465 |
|  | ±4ms | 385 | 1449 | 2226 | 4059 | 5.0 | 39.4 | 81.9 | 89.9 | 24.5 | 250 | 518 | 533 |
|  | ±6ms | 385 | 1450 | 2226 | 4061 | 6.5 | 44.9 | 93.8 | 101 | 38.8 | 287 | 560 | 624 |
| Method 3 | ±2ms | 385 | 1448 | 2218 | 4051 | 3.5 | 28.9 | 72.0 | 75.8 | 21.1 | 165 | 434 | 462 |
|  | ±4ms | 385 | 1448 | 2216 | 4049 | 4.4 | 35.2 | 79.2 | 79.7 | 25.3 | 195 | 568 | 472 |
|  | ±6ms | 385 | 1447 | 2223 | 4057 | 5.5 | 44.0 | 98.1 | 103 | 34.4 | 274 | 603 | 687 |
| Method 4 | ±2ms | 385 | 1449 | 2218 | 4052 | 3.3 | 28.8 | 70.1 | 73.1 | 19.7 | 160 | 431 | 487 |
|  | ±4ms | 385 | 1451 | 2217 | 4051 | 4.5 | 40.1 | 79.1 | 88.9 | 25.7 | 257 | 508 | 599 |
|  | ±6ms | 385 | 1458 | 2224 | 4064 | 6.1 | 56.0 | 110 | 129 | 31.6 | 397 | 714 | 850 |
|  |  | maxF [mN] |  |  |  | mean freq [Hz] |  |  |  |  |  |  |  |
|  | NS | 393 | 1519 | 2398 | 4234 | 13.6 | 24.0 | 36.1 | 35.6 |  |  |  |  |
| Method 1 | ±2ms | 395 | 1523 | 2406 | 4239 | 12.6 | 23.8 | 35.7 | 34.1 |  |  |  |  |
|  | ±4ms | 399 | 1557 | 2457 | 4357 | 11.3 | 22.6 | 33.1 | 31.5 |  |  |  |  |
|  | ±6ms | 402 | 1627 | 2460 | 4370 | 10.2 | 22.7 | 32.2 | 31.5 |  |  |  |  |
| Method 2 | ±2ms | 394 | 1517 | 2407 | 4258 | 12.9 | 24.0 | 35.7 | 34.2 |  |  |  |  |
|  | ±4ms | 397 | 1586 | 2456 | 4316 | 10.9 | 20.7 | 33.1 | 30.9 |  |  |  |  |
|  | ±6ms | 402 | 1588 | 2482 | 4367 | 10.4 | 23.6 | 29.8 | 30.0 |  |  |  |  |
| Method 3 | ±2ms | 395 | 1531 | 2407 | 4259 | 13.3 | 24.1 | 35.7 | 34.4 |  |  |  |  |
|  | ±4ms | 399 | 1544 | 2462 | 4256 | 12.1 | 22.4 | 33.2 | 33.9 |  |  |  |  |
|  | ±6ms | 404 | 1574 | 2517 | 4330 | 11.5 | 21.6 | 30.8 | 31.2 |  |  |  |  |

|  |  |  |  |  |  |  |  |  |  |
| --- | --- | --- | --- | --- | --- | --- | --- | --- | --- |
| Method<br>4 | $\pm 2\text{ms}$ | 394 | 1523 | 2399 | 4250 | 13.4 | 23.8 | 35.5 | 34.6 |
| | $\pm 4\text{ms}$ | 398 | 1587 | 2472 | 4331 | 10.8 | 21.2 | 32.3 | 29.5 |
| | $\pm 6\text{ms}$ | 399 | 1626 | 2565 | 4440 | 10.8 | 21.6 | 25.9 | 24.4 |

- *Force mean value*: The synchronization had no effect on *meanF* value, showing a negligible change (≤ 12 mN) before and, after the synchronization was applied, i.e., for the muscle force, *meanF* was varying from 4052 mN (*NS*) to a maximum of 4064 mN (Method 4, Δt = ± 6 ms) (Table 2). This can be also tracked in Fig. 7A, which presents no visible difference in the baseline value (red solid horizontal line) when comparing all forces placed in a row.
- *Force RMS value*: The synchronization had an important effect on increasing the *rmsF* value by more than 50 mN, which could become as high as 129 mN for the muscle force (Method 4, Δt = ± 6 ms), considering its baseline *NS* value of 73.5 mN (Table 2). Additionally, Fig. 8B is provided to show the relative *rmsF* change as a ratio of synchronized vs. *NS* value. Specifically, it shows that the maximal *rmsF* increment (about two times) could be achieved for the forces of two types of MUs (S, FR) following synchronization with Methods 1, 2, and 4, Δt = ± 6 ms. Considering the whole muscle, the observed maximal increment of *rmsF* was about 1.8 times, achieved using Method 4, Δt = ± 6 ms.
- *Force min–max range*: The synchronization had an important effect on increasing the *rangeF* value by about 450 mN, which increased from 405 mN (NS) up to 850 mN for the muscle force after synchronization with Method 4, Δt = ± 6 ms (Table 2). The *rangeF* ratio (synchronized vs. *NS* value) in Fig. 8C shows that the largest *rangeF* increment (two to 2.6 times) was achieved for the forces of two types of MUs (S, FR) after synchronization with Methods 1, 2 and 4, Δt = ± 6 ms. Considering the whole muscle, the observed maximal increment of *rangeF* (about 2.1 times) was with Method 4, Δt = ± 6 ms. Although the observations concerning *rangeF* are similar to those of *rmsF* as was noted above, the relative and absolute changes in *rangeF* values as an effect of synchronization were larger. This could also be visually confirmed by the force signals in Fig. 7A (blue dotted lines show larger span than red dotted lines after synchronization, comparing all forces placed in a row).
- *Maximal force*: The synchronization had an important effect on increasing the *maxF* value by more than 205 mN, which could raise it from 4234 mN (*NS*) up to 4440 mN for the muscle force after synchronization with Method 4, Δt = ± 6 ms (Table 2). The *maxF* ratio (synchronized vs. *NS* value) in Fig. 8D shows that the largest *maxF* increment (up to 1.7 times) is achieved for the forces of two types of MUs (FR, FF) after synchronization with Method 4 or Method 1, Δt = ± 6 ms. Considering the whole muscle, the observed maximal increment of *maxF* was about 1.5 times using Method 4, Δt = ± 6 ms. This relative change of *maxF* was found to be smaller than the force amplitude variations estimated above by the other two force parameters (*rangeF* and *rmsF*).
- *Force mean spectral frequency:* In this context, the synchronization had an important effect—decreasing the *meanfreq* value by more than 10 Hz, which drops it from 35.6 Hz (*NS*) down to 24.4 Hz for the muscle force after synchronization with Method 4, Δt = ± 6 ms (Table 2). The *meanfreq* ratio (synchronized vs. *NS* value) in Fig. 8E shows that the largest *meanfreq* drop (i.e., < 0.75 or > 25% vs. *NS*) could be achieved for the forces of two types of MUs (S, FF) after synchronization with Methods 1, 2 and 4, Δt = ± 6 ms. Considering the whole muscle, the observed maximum drop of *meanfreq* was about 30% (< 0.7 Hz) with Method 4, Δt = ± 6 ms. This can be observed in the *PSD* of Fig. 7B (first row for the muscle force and second row for FF MU force) as a shift of the high-frequency components (predominantly around 40 Hz) in the *NS* spectrum to low-frequency components (10–25 Hz) in the spectrum for synchronization with Method 4, Δt = ± 6 ms.

### Correlation of the force variance and MU synchronization

The results presented in this section aim to answer the general question of whether the provided synchronization methods regularized by widening the time window for synchronization (Δt = ± 2, ± 4, and ± 6 ms) led to consistent increases in both the level of MU synchronization (*CISI*) and the induced changes in the force output. Thus, the force parameters, which were most closely correlated to the synchronization design in Methods 1 through 4, could be deduced. The results in Table 3 establish the correlations between the curves in Fig. 8A for the level of MU synchronization in the function of Δt (*CISI* = *f*(Δt)) and each of the curves in Figs. 8B through 8E for the variance of the five force parameters as a function of Δt (*meanF, rmsF, rangeF, maxF, meanfreq*). The correlations were estimated in the range [−1;+1], where +1 and −1 stand for strongly correlated curves that were directly or inversely proportional, respectively. The results show that *rmsF, rangeF* and *maxF* were the most robust force parameters, which were consistently increased by the synchronization level with an average correlation coefficient of 0.97; the force mean spectral frequency was indeed inversely proportional to the synchronization level, with an average correlation coefficient of −0.89; and *meanF* was the parameter least dependent on the synchronization, with an average correlation coefficient of 0.53.

**Table 3.**
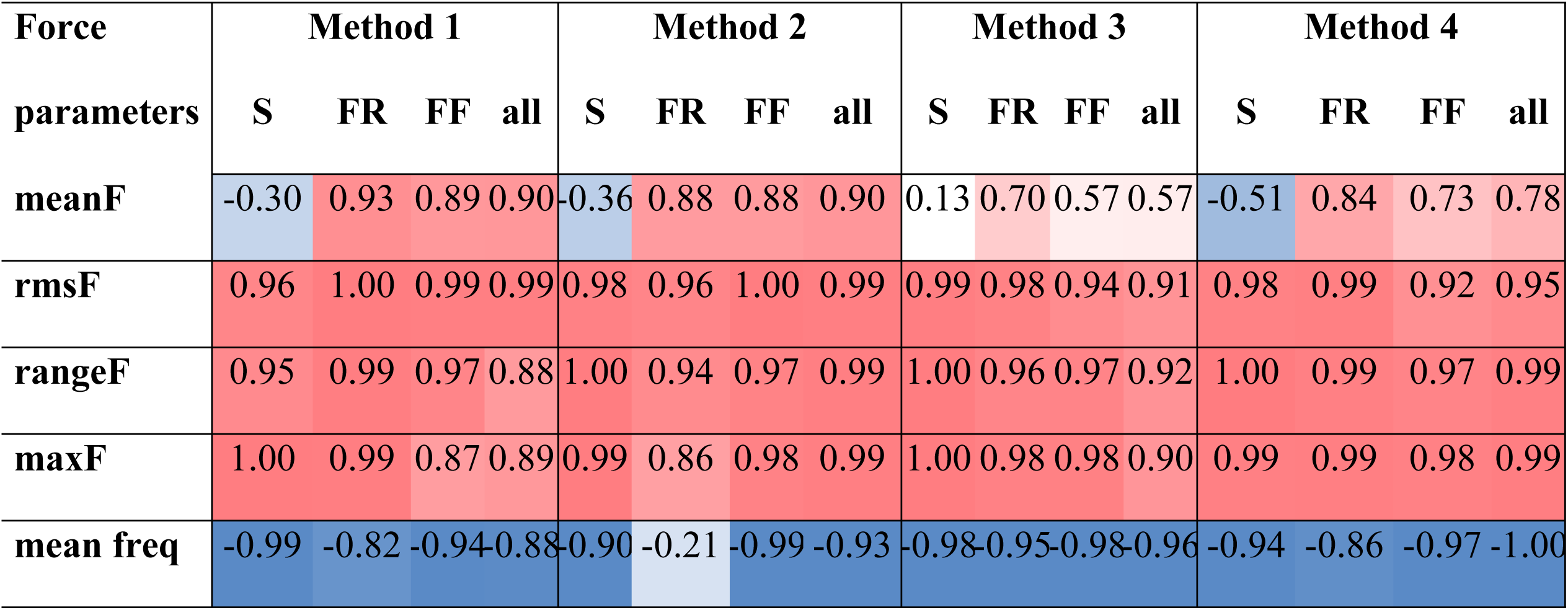
Correlation coefficients between *CISI* and the five force parameters *meanF, rmsF, rangeF, maxF*, and *meanfreq*. The strength of the correlation is coded with a color gradient, highlighting the strong positive (> 0.8) (dense red) and strong negative (< −0.8) (dense blue) correlations.

## Discussion

There are two different approaches one could use to investigate the synchrony between different MUs and its influence on the developed muscle force. The first one involves assessing experimental recordings of electromyographic signals using needle or surface electrodes and decomposing these signals into individual action potentials [4, 37-40]. However, the disadvantages of this approach include that only a portion of the active MUs is recorded, it is not possible to distinguish fast from slow MUs and the measured muscle force reflects the force of all active MUs, and even MUs of other muscles. The second method is based on pure modeling, wherein models of the muscle are composed using different MUs [21, 31]. These models are based on the Fuglevand et al. approach [30] and contain 120 MUs. However, these authors did not divide MUs into different types (S, FR, and FF). Moreover, the function used for describing the twitch was based only on two parameters: force amplitude and contraction time. The model used in the current paper, constructed based on experimental data concerning MU twitch and tetanus properties as well as motoneuronal excitability, has been fully described previously [33]. Here, the experimentally measured twitches are modeled by a six-parameter function and the summation of the twitches into tetanus is established by an experimentally verified mathematical algorithm. In the adopted basic model, it was proven that the firing of all MUs is asynchronous. Then, synchronization was imposed in this basic MU firing arrangement, changing the pattern of impulses of MUs during the steady state of the muscle force using several simulated situations (i.e., four modes of synchronization with the three time windows ± 2, ± 4, and ± 6 ms). In this way, broad investigation of the influence of the synchrony of the three types of the MUs on the developed muscle force and cumulative forces of MUs from the three groups could be performed. The results based on the two used coefficients *corMU* and *CISI* showed that the range, the maximum, and the root-mean-square of the forces rose with increased synchronization, while the mean forces remained nearly unchanged. This increase was stronger for fast MUs; notably, these units are mostly responsible for the force instability (muscle tremor) in the context of moderate or strong muscle contractions, wherein fast MUs are recruited into activity.

### Models of MU synchronization

To increase the degree of synchronization and to analyze its effects on the muscle and MU forces, we considered the synchronization of pulses of pairs of MUs in the time windows ± 2, ± 4, and ± 6 ms. It is known that synchronization is an effect of a common excitatory input to several motoneurons and that synchronic excitatory postsynaptic potentials (EPSPs) evoked in several motoneurons increase the probability of the simultaneous occurrence of their action potentials [41]. The size of the time windows is related to the duration of EPSPs in rat motoneurons, lasting several milliseconds, with an increasing phase often remaining below 2 ms (for example, for Ia monosynaptic EPSPs, see Fig. 1 in Seburn and Cope [42]. Additionally, the applied method resulted in a narrow peak in the cross-interval histogram (Fig. 6), similar to that reported for human muscles by De Luca et al. [4], as is typical for short-term synchronization (i.e., the peak centered about zero-time delay 0.5 ± 2.9 ms) and with an average width of 4.5 ± 2.5 ms. For all four proposed modes of synchronization used in the present study, the same range of time windows was applied. The largest (± 6 ms) time window increased the *CISI* by about 1.5 times for Method 1, about 2.5 times for Methods 2 and 3, and more than three times for Method 4 (see Table 1). The range of differences in the obtained synchronization is similar to that of differences in the *CISI* reported for trained and nontrained subjects (more than two times higher in weightlifters), changes resulting from conditioning exercise (about 2.5 times higher after the exercise), and those between dominant and nondominant hands (1.6 times higher in the nondominant hand) [39].

The proposed method of inducing synchronization within time windows Δt of variable duration appeared to be an efficient tool in the four tested simulations. For all four methods of synchronization, values of the investigated parameters *musF, rangeF*, and *maxF*, which characterized the force oscillations, rose along with increases in the time window Δt, i.e., when the synchronization degree was augmented (Fig. 8). Notably, this increase appeared strongest with Method 4 and weakest with Method 3. Meanwhile, the highest value of *corMU* (74.6) was achieved for Δt = ± 6 ms for FR MUs (Table 1). Moreover, except for in Method 3, the highest values of *corMU* were observed for FR MUs (Table 1). This observation is surprising in light of previous physiological experiments concerning force decreases/increases resulting from the prolongation/shortening of one IPI during the unfused tetanic contraction ascertained using MUs of the rat medial gastrocnemius [28]. Namely, relative force fluctuations noted for FF and FR MUs were similar and one could expect no differences to exist between these two types of fast MUs in the present simulation study. This methodological approach resulted in the highest synchronization for Method 4 and is reflected by the parameters *corMU* and *CISI* in Table 1. It should be stressed that the four methods led to similar effects on muscle force—that is, greater maximal force and higher fluctuations around a mean force—and these increases concerned all three types of MUs, although it should be stressed that this result was obtained for the maximum excitation signal, i.e., a simulation of a very strong contraction, when all MUs were active.

### Effects of synchronization on MU and muscle forces

The influence of the increasing synchronization level on the mean as well as on the maximum force of particular MU types and of the whole muscle was, in general, very weak (i.e., the maximum force increased by up to 5% for the whole muscle and up to 7% for FF MUs), regardless of the synchronization method applied in the model. This confirms the results of previous studies, which also demonstrated that the magnitude of force output and the average force of the muscle were not altered considerably due to synchronization [21, 39]. However, an increase in the synchronization time window from ± 2 to ± 6 ms in all cases correlated with a rise in the force of each MU type, with the change being the greatest for synchronization Methods 1 and 4. Moreover, the present study has revealed certain differences between MU types. Not only did the absolute force increase but also the relative force increased after synchronization; further, they were always the highest for fast MUs (FF and FR) and the lowest for slow MUs. This also confirms previous observations that synchronization may be beneficial during the performance of contractions where rapid force development is required, for which fast MUs should be recruited [17].

On the other hand, it was already mentioned that a muscle can produce smooth contractions due to asynchronous discharges of motor neurons [17, 23] and that synchronization increases the variability in the muscle force [21]. Indeed, simulated contractions in our model have confirmed that synchronization substantially influences the range of force oscillations during the steady state of the muscle contraction and the min–max range of modeled forces gradually rose with the increase in the time window for synchronization in each method. This can be partly explained by previous computer simulations indicating that synchronization leads to an increase in the estimated twitch force and to a decrease in the estimated contraction time of an MU [26]. Obviously, absolute values of the min–max range of the force were the lowest for the weakest S MUs, but the ratio of the *rangeF* parameters (as well as the ratio of the *rmsF* parameters) between synchronized and *NS* models was the highest for S MUs for all methods—except Method 4, in which MUs of the same type were synchronized according to the first MU in the group (see Figs. 8B and 8C).

A 100% excitation signal (corresponding to a very strong muscle contraction) used in this model was applied to ensure activation of all MU types, which helped us to elucidate the contributions of the three types of MUs to muscle tremor, which are dependent on the force level [43]. According to the size principle, at a lower excitation signal, a contribution of high-threshold fast MUs (especially those of the FF type) to the force development would be smaller or recruitment would be restricted to low-threshold (S or S and FR) MUs. The lowest relative force oscillations were noted in FF MUs for all methods of synchronization (Fig. 8B). This observation indicates that slow MUs have the strongest and FF MUs have the weakest relative influence, respectively, on force fluctuations described as muscle tremor and thus partly explains why tremor is best visible during weak contractions, when predominantly slow MUs are recruited.

### The influence of synchronization on the spectral frequency of the muscle force

To our knowledge, the parameter *meanfreq* of the force has not been analyzed in muscle modeling in connection with the synchronization of MU firing to date. It should be noted, however, that the power spectral analysis of tremor in the first dorsal interosseous muscle revealed three frequency peaks occurring at around 10 Hz, 20 Hz, and 40 Hz [24], which correspond to our findings concerning the mean spectral frequencies of S, FR, and FF MUs, respectively (Table 2). McAuley et al. [24] concluded that their results reflected the synchronization of MUs at frequencies determined by oscillations within the central nervous system; however, our findings suggest that the force oscillations related to three types of MUs likely contribute to those frequency peaks.

A decrease in the *meanfreq* was observed in parallel with an increase in the degree of synchronization in all four applied methods. It should be stressed that the mean firing frequencies of all MUs remained unchanged during simulations, due to a constant number of pulses in the analyzed time window (2000 ms). A decrease in force spectral frequencies paired with the occurrence of slower force oscillations. This observation at increased synchronization levels indicates that the force-frequency spectrum depends upon the temporal distribution rather than on the mean firing frequencies of MUs and this conclusion concerns all three types of MUs, despite considerable differences in the *meanfreq* between S, FR, and FF MUs. The decrease in the *meanfreq* could not be linked to muscle fatigue, which was not modeled, and should instead be connected with processes of summation of twitches into tetanic contractions.

McAuley and Marsden [44] in their review argued that the physiological tremor in humans is likely of multifactorial origin, with contributions from the 10-Hz range of oscillatory activity of the central nervous system, MU discharge frequencies, reflex loop resonances, and mechanical resonances. However, it must be emphasized that the present results were obtained using the model of a rat muscle, so it is risky to directly compare the frequencies related to different types of MUs collected herein to human data, most of all because rat MUs demonstrate considerably faster contractions and have higher discharge frequencies.

### Conclusion

The present study revealed that, regardless of the method used for the synchronization of MU firings, the increase in the synchronization index had a negligible effect on the mean force of the developed contractions yet influenced muscle tremor by increasing force oscillations and further highlighted that these results were observed for all three types of MUs. A parallel decrease in the mean spectral frequency of the force indicated that, in the synchronized models, the force oscillations were slower despite higher magnitudes. The synchronization of fast MUs led to higher increases in the range of the force variability and the force root-mean-square in comparison with that of slow MUs. On the other hand, relative changes in the latter parameters in the synchronized simulations were the highest for slow MUs, indicating their significant contribution to muscle tremor, especially during weak contractions.

## Acknowledgments

The study was supported by a bilateral agreement between the Bulgarian Academy of Sciences and the Polish Academy of Sciences.

## Author contributions

Conceptualization: RR, JC, PK

Data curation: RR, VK

Formal analysis: RR, VK, PK, JC

Funding acquisition: RR, JC

Investigation: RR, VK, PK, HDC, KK, JC

Project administration: RR, JC

Software: RR, VK

Validation: RR, VK, PK

Writing—original draft: RR, VK, PK, JC, KK

Writing—review and editing: RR, VK, PK, HDC, KK, JC

